# Roles of motor on-rate and cargo mobility in intracellular transport

**DOI:** 10.1101/2020.07.13.201434

**Authors:** Matthew J. Bovyn, Babu J.N. Reddy, Steven P. Gross, Jun F. Allard

## Abstract

Molecular motors like kinesin are critical for cellular organization and biological function including in neurons. There is detailed understanding of how they move and how factors such as applied force and the presence of microtubule-associated proteins can alter this single-motor travel. In order to walk, the cargo-motor complex must first attach to a microtubule. This attachment process is less studied. Here, we use a combination of single-molecule bead experiments, modeling, and simulation to examine how cargos with kinesin-1 bind to microtubules. In experiment, we find that increasing cargo size and environment viscosity both signficantly slow cargo binding time. We use modeling and simulation to examine how the single motor on rate translates to the on rate of the cargo. Combining experiment and modeling allows us to estimate the single motor on rate as 100 s^−1^. This is a much higher value than previous estimates. We attribute the difference between our measurements and previous estimates to two factors: first, we are directly measuring initial motor attachment (as opposed to re-binding of a second motor) and second, the theoretical framework allows us to account for missed events (i.e. binding events not detected by the experiments due to their short duration). This indicates that the mobility of the cargo itself, determined by its size and interaction with the cytoplasmic environment, play a previously underestimated role in determining intracellular transport kinetics.

## 1 Introduction

Eukaryotic cells employ molecular motors to create and maintain order, in part via transporting organelles from one place in the cell to another. At a single-molecule level, motor behavior involves attachment to the microtubule, stepping, and detachment from the microtubule. While the stepping and detachment processes have been extensively studied — including under load in backward, forward [1], left, right [2], and upward [3] directions, and as modified by MAPs [4, 5] and tubulin state [6] — the attachment rate is less well-understood.

Although rate constants can be used to summarize the attachment process, the study of motor attachment is complicated by multiple notions of on-rate: First, there is the “landing rate” of motors from solution onto an immobilized microtubule. Second, there is the overall time for a cargo adorned with motors to attach to a nearby microtubule. This cargo attachment process involves the subprocess of an individual motor attaching to the microtubule, corresponding to yet a third on-rate (which we refer to as *k*_on_). Fourth and finally, for cargo with multiple motors, it is possible that once one motor is attached, subsequent motors bind at a different rate than the first. These different notions of rates of kinesin can be inferred from several studies. Landing rate assays (e.g. [7]) measure binding by directly detecting events where individual motors bind the microtubule. Because binding is detected directly, little interpretation of the data is necessary quantify the landing rate. However, because the motors are not bound to a cargo, the landing rate is difficult to translate into the context of a motor bound to a cargo. Force generation and run length of systems where two or more motors work together have been measured when linked directly to the bead [4, 8], on antibody linkers attached to beads [9], on DNA linkers without beads [10, 11], and on DNA linkers with beads [12, 13]. Because the motors are necessarily close together in these constructs, the bound or unbound state of each motor is difficult to detect directly (with the notable exception of [14]). Instead, mathematical models have been used to interpret motor binding rate by fitting model parameters to data. The on rates gleaned from these experiments differ from landing rates in two ways. First, the motors are linked together with beads and other molecules, so where the motors are relative to each other and how they are able to move through space is different in each case. Second, the on rate measured is always the rate at which the second motor binds (or perhaps third or fourth, depending on the experiment). The on-rate for kinesin of around 5 s^−1^ was estimated by fitting an ODE model to data on nanotube extraction in [15] and for multiple motor transport on beads in [8]. In an experiment where motor number was controlled using antibody anchoring points, motor on rate was estimated as 0.7 s^−1^ [9]. To match run lengths of multiple motor motor beads, 10 s^−1^ was used in previous work by us [16]. For a two-motor ensemble constructed with a DNA linkage but no cargo bead, a motor on rate of 4.7 s^−1^ was found [14].

Here we use optical tweezers on a bead assay, along with biophysical modeling, to investigate the full process of cargo binding to a nearby microtubule. By analyzing the bead assay time course data, we are able to extract times for the cargo to bind the microtubule and correct for binding events that are missed because they are hidden by thermal fluctuations. We perform experiments on beads of varying sizes and in mediums of varying viscosity. Putting this data together suggests that binding time is heavily dependent on bead mobility, specifically its rotation. To understand the relationship between binding time and bead rotation, we construct a biophysical model and find that this model recapitulates the experimentally derived dependance of binding time on cargo rotation. We use the bead binding data to estimate parameters of the model. We find *k*_on_ ≈ 100 s^−1^, which is considerably higher than previous estimates. This fast motor on-rate implies a greater importance for the timescales associated with cargo mobility in determining intracellular transport.

## 2 Results

### 2.1 Bead assay and simulations together reveal cargo rebinding times

To investigate the binding of cargos to microtubules, we use a bead assay [4, 16, 17] altered to ensure the center of the trap is near the microtubule surface, described in methods section 4.1. We construct in vitro cargos with a single motor, and use an optical trap to bring the cargo near the microtubule. We monitor the cargo position as a function of time (figure 1A, lower): as the motor binds the microtubule and subsequently walks, it displaces the cargo in the trap (figure 1Ai). Because we use an optical trap stronger than the motor’s stall force, the motor always either stalls or detaches, and eventually falls back to the trap center (figure 1Aii), rather than escaping and continuing along the microtubule. There is a period of time after the bead falls back to the center of the trap during which we observe only small fluctuations in bead position (figure 1Aiii). Eventually the motor rebinds, and displaces the bead in the trap once more (figure 1Av). We monitor beads for several minutes, during which this cycle repeats 10-100 times. For the motor to move the bead after detachment, it must first bind to the microtubule. This must happen some time after the bead starts to fall back to the trap center, and before movement of the bead is detected again (figure 1Avi).

**Figure 1:**
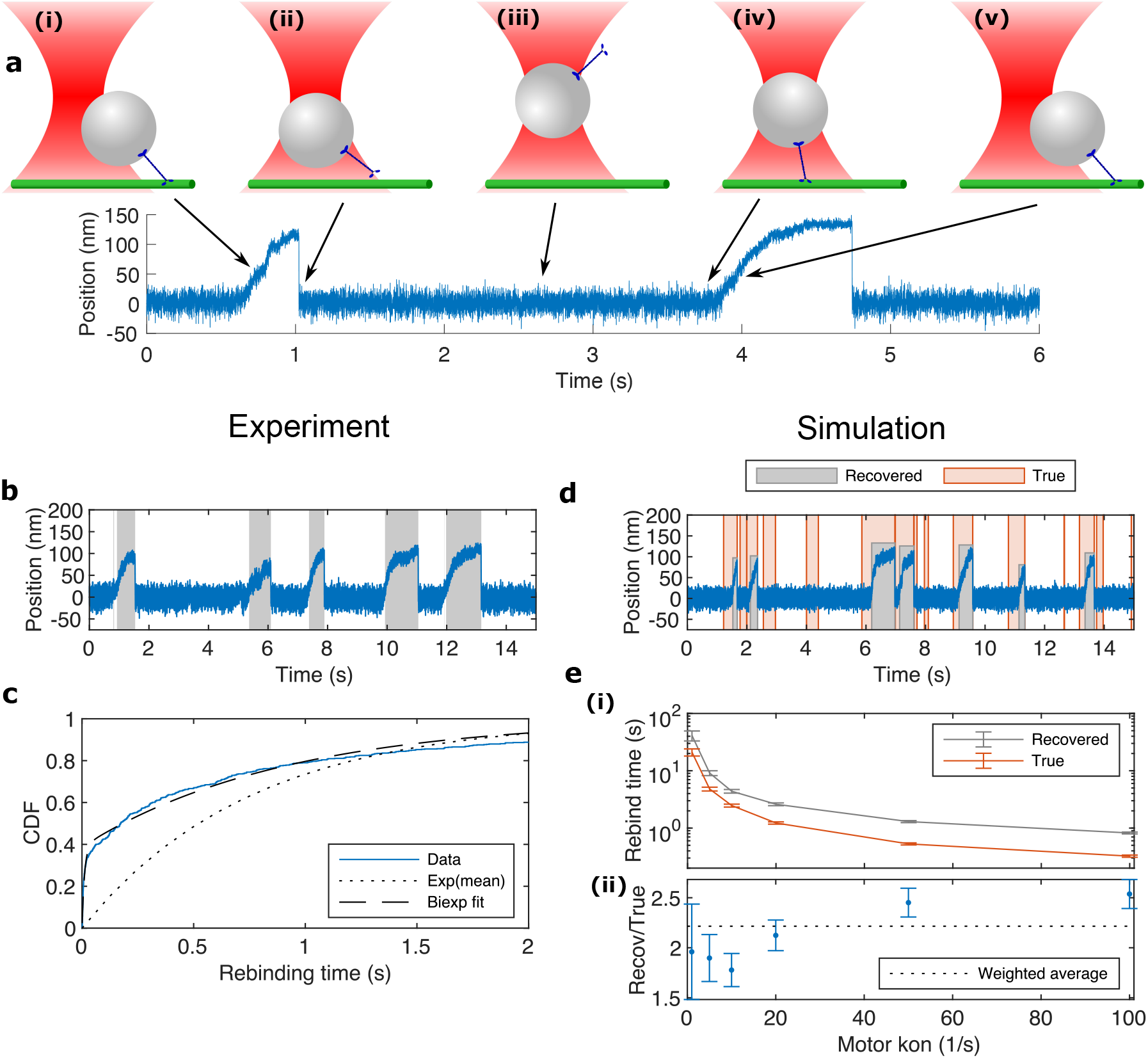
Experiments and simulations together reveal cargo rebinding time. a) In our bead assay, we record the position of the cargo with time (bottom). We observe the bead pull the cargo out from the trap center (i). The motor detaches from the microtubule and the cargo falls back to the trap center (ii). After this, the bead diffuses in the trap (iii) until the motor binds the microtubule again (iv) and we are able to observe it move the bead (v). b) We detect when the bead is pulled from the center of the trap using custom software. For the raw trace in blue, the recovered periods of time where the motor is bound are highlighted in grey. c) Using the labeled time series, we extract rebinding times. The empirical cumulative distribution function is shown along with exponential and biexponential fits. d) Using simulation, we generate traces of a single-motor bead in a trap. We compare the true bound state of the motor from simulation (orange highlighting) to the bound state recovered by the algorithm (grey highlighting). e) Comparison of the true rebinding time to the one recovered by the algorithm. We show a range of motor on rates, which result in rebinding times from more than 10 s to less than 1 s (i). Over this range, the recovered rebinding time is always greater than the true rebinding time. Quantitatively, we find the recovered rebinding time is on average 2.2 times the true rebinding time (ii). Lines between points in (i) are guides for the eye. Error bars are standard error of the mean.

From the bead position data, we algorithmically mark the start and end times of binding events (see methods section 4.2). We define the rebinding time as the time between detection of an unbinding event and the next detection of bead movement. An experimental trace with highlighted binding events is shown in figure 1B. We extract rebinding times from this annotated data. Rebinding times for 560 nm diameter beads have a mean of 0.80 ± 0.05 s (mean ± SEM). The distribution is shown in figure 1C. The distribution of rebinding times differs from an exponential distribution of the same mean by having more short events, and is better fit by a biexponential. This suggests the binding process is more complex than a single rate process.

To understand how the detected binding events relate to the true bound state of the motor, we next used a cargo transport simulator constructed previously [16] to generate simulated traces. This simulator explicitly includes the diffusion of a small sphere in a fluid as well as motor on-off and stepping dynamics. It calculates the 3D translational and rotational motion of the bead resulting from forces exerted by the optical trap, motor stepping, and diffusion. Rates for motor binding, unbinding and stepping depend on instantaneous spatial position and force on the motor. For more details, see methods section 4.5.

The simulated position traces are qualitatively similar to the experimental ones (figure 1d). We run our simulated traces through the kick-finding algorithm used on the experimental data. We find that, as expected, the motor often binds a significant time before the displacement of the bead is detectable in the position trace, as shown in figure 1d. Unexpectedly, we also find a significant number of binding events where deflection of the bead is undetectable in the position trace. When we visualize these events in the simulation, we find that the motor binds and begins to walk, but then unbinds before it has walked far enough to begin pulling the bead out of the trap. This is demonstrated in supplemental video 1. The distribution of rebinding times from the simulation, shown in S2, is also consistent with the fraction of hidden events and bi-exponential distribution.

The simulation takes into account subtleties in interpreting the attachment process. For example, the amount of time after motor binding but before detection of bead movement may vary. If the bead binds in such a way that the motor is closer to the plus end of the microtubule than the bead center, the motor has few steps to take before it begins displacing the bead. However, if the motor binds such that the motor heads are closer to the minus end of the microtubule than the bead center, the motor may walk farther than a bead diameter before it begins displacing the bead in the trap. Walking this distance takes our kinesin motors half a second or more.

We then compare how these detection effects change the calculated rebinding time. In figure 1e, we show that the recovered rebinding time is always longer than the true rebinding time in simulation. Over a large range of rebinding times, the recovered rebinding time is consistently about twice the true time.

### 2.2 Time to bind depends on cargo and environment

Having established that we can recover and interpret cargo rebinding times, we next asked how rebinding time depends on the cargo size and the environment around it.

To assess the impact of properties of the environment on rebinding time, we added sucrose to the experimental medium. Adding sucrose increases the viscosity of the fluid [18]. We find that higher concentrations of sucrose yield longer rebinding times, as shown in figure 2A. When enough sucrose is added to increase the viscosity of the experimental medium by 10 times, the mean rebinding time increases by a factor of 5. Therefore rebinding time for a cargo depends on the environment of the cargo.

**Figure 2:**
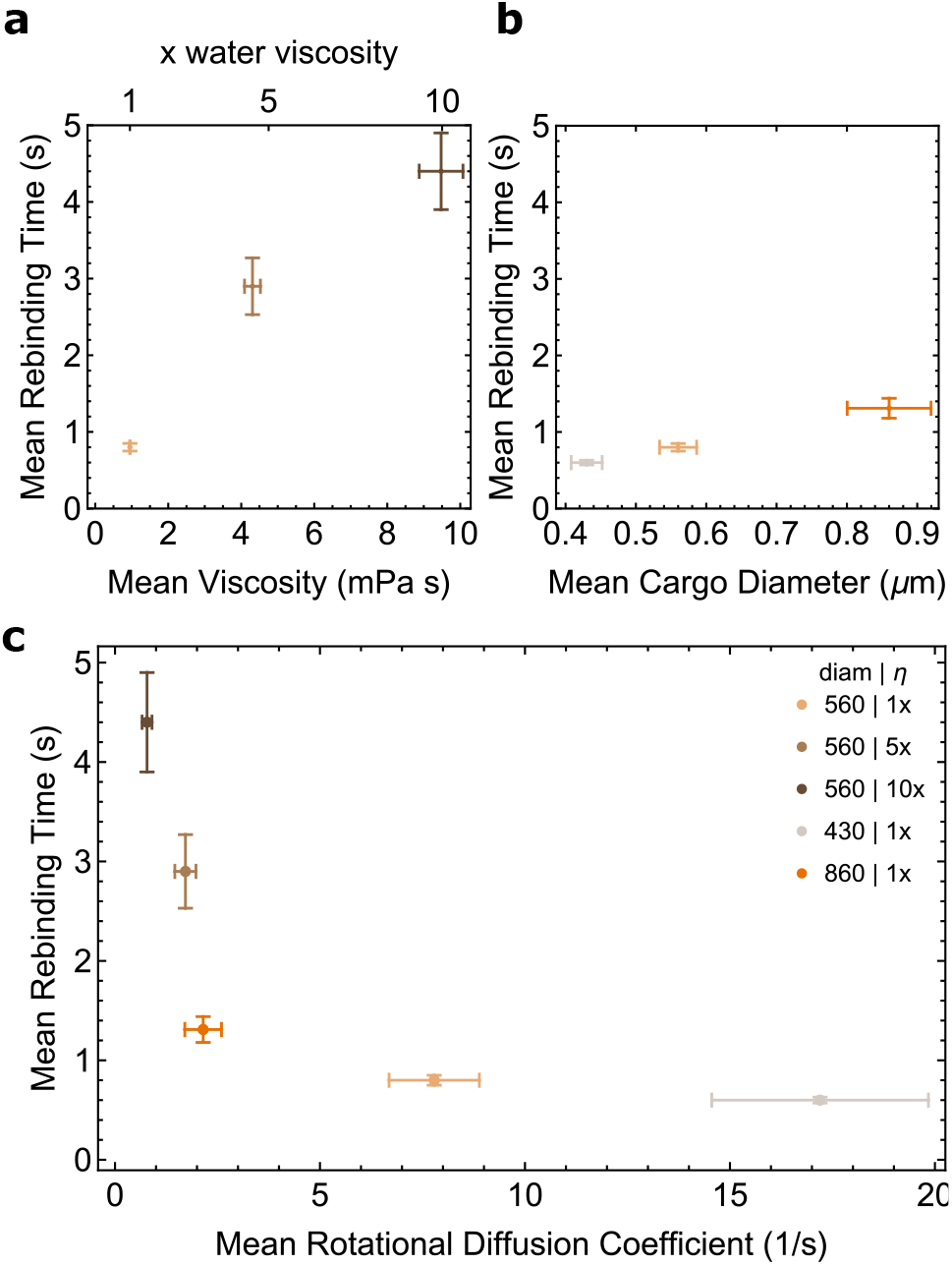
Measured rebinding times at different viscosities and bead diameters. Colors in all panels correspond to legend in panel c. a) Measured rebinding times in buffer with sucrose added to increase viscosity. b) Measured rebinding time for cargos of different diameters. No sucrose was added to the buffer. c) Data from a and b plotted on against rotational diffusion coefficient of the bead, *D* = *k_B_T*/(8*πηR*^3^), where *η* is viscosity and *R* is radius. All errorbars are standard error of the mean. Measurements reflect about 10 beads measured, with dozens of rebinding events for each bead. For more information on error estimation, see methods.

To asses how properties of the cargo impact rebinding time, we measured rebinding times for cargos of a variety of diameters. We find that larger cargos take longer to rebind on average, as shown in figure 2B. Mean rebinding time for 860 nm diameter cargos is about double that for 440 nm cargos. Therefore rebinding time depends on properties of the cargo itself, in particular its size.

Experimental findings for all conditions are summarized in table 1.

**Table 1:** Rebinding times measured for beads of different sizes and in solutions of different viscosities.

|  | 430 nm | 560 nm | 860 nm |
| --- | --- | --- | --- |
| 1x | 0.6±.03 | 0.8±.05 | 1.3±.13 |
| 5x |  | 2.9±.37 |  |
| 10x |  | 4.4±.50 |  |

Because changes in the translation and rotation dynamics of the bead are common to both experimental perturbations, we ask if these effects might be able to explain the changes in rebinding time. Thermal physics describes that the movement of a small isolated sphere in a viscous medium is governed by diffusion equations. Movement speed is proportional to the diffusion coefficient, which for rotation is *D* = *k*_*B*_*T*/(8*πηR*^3^), where *k_B_* is Boltzmann constant, *T* is the temperature of the system, *η* is the viscosity of the medium and *R* is the radius. We plot all the experimental data as a function of the rotational diffusion coefficient for that condition in figure 2. We find that the data appears to collapse onto a single curve, suggesting that changes to bead rotation dynamics are the driving factor in changes to the rebinding time in the experiments.

### 2.3 A compartmental model reveals dependance of rebinding time behavior on cargo size and viscosity

To understand if changes to cargo rotation—and hence ability of the cargo-bound motor to reach the microtubule—can explain the changes to the rebinding times of the beads in the experiments, we a construct hybrid continuous time Markov chain model with rates constrained by the detailed simulation. This model assigns the system to be in one of three states. First, the cargo can be in a configuration where the motor heads cannot reach the microtubule e.g. because the bead is in its way. We call this state “far”. Otherwise, the state is “near”. Finally, the motor may be bound to the microtubule. We call this state “engaged”. States and possible transitions between them are shown in figure 3A.

**Figure 3:**
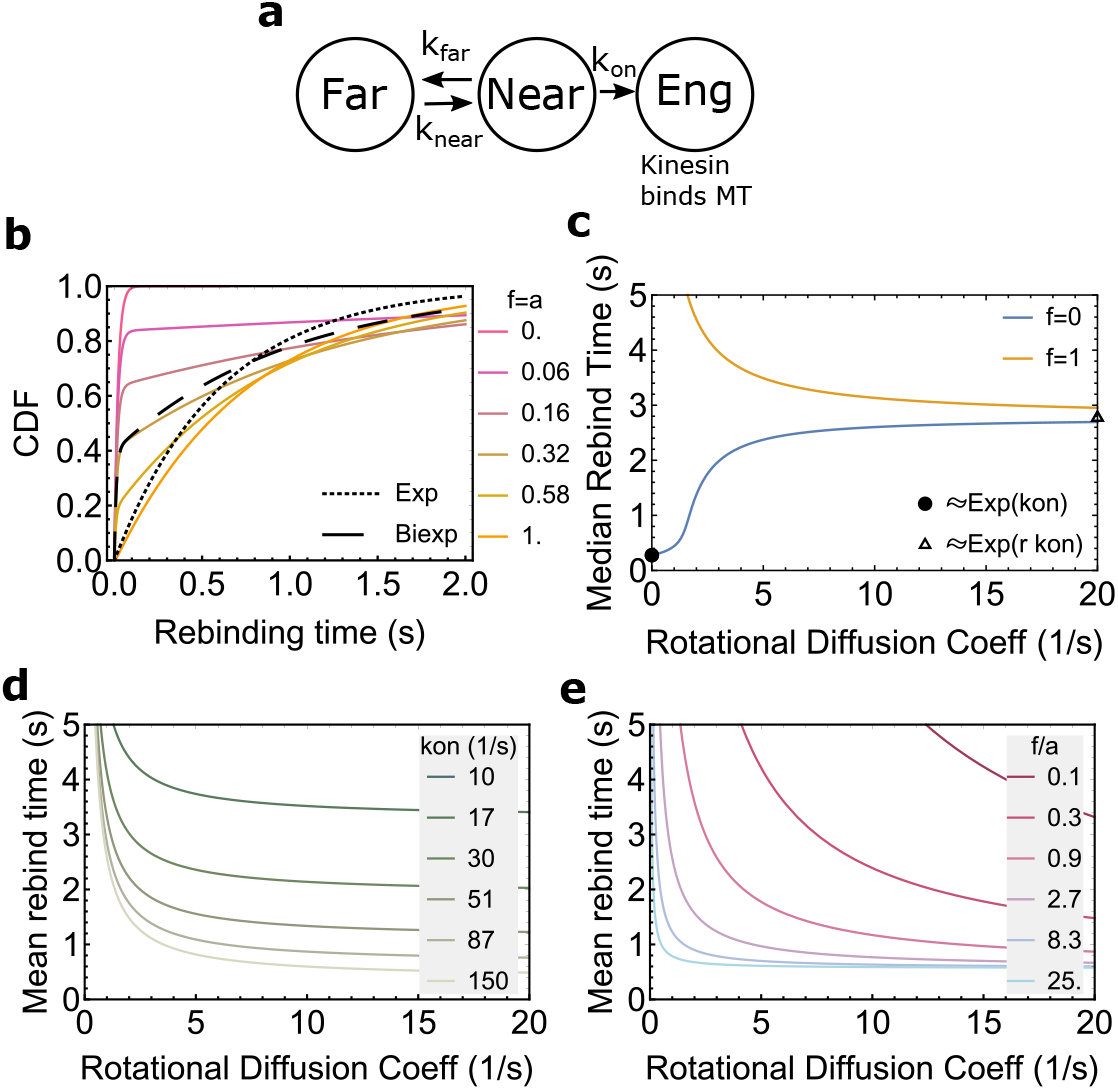
Model for rebinding time We construct a Markov chain model to describe rebinding. It has 3 states: motor far from the microtubule, motor within binding range of the microtubule, and motor engaged on the microtubule (a). The distribution of rebinding times resulting from the model (b) can exhibit both exponential (near *f* = *a* = 1 or *f* = *a* = 0) and bi-exponential behavior (*f* = *a* between 0 and 1). Near *f* = *a* = .2, the model gives a close match to the bi-exponential fit to the experimental rebinding distribution for 560 nm diameter beads in water (dashed curve). We calculate the mean time to enter the engaged state. Because of a quirk of the mean when combined with rare events, this expression is not well behaved when *f* = 0 and *D* is small. We therefore examine the median instead in (c). At high rotational diffusion coefficient, the rebinding time goes 1/(*rk*_on_), where r is set by the geometry of the bead and motor. At low rotational diffusion coefficient, the behavior of the model depends on the parameter *f*/*a*. For *f*/*a* = 0, rebinding time goes to 1/*k*_on_. For *f*/*a* > 0, time goes to infinity. Behavior between these two extremes is dictated by a combination of parameters *k*_on_ and *f*/*a*, as show in (d) and (e).

To understand how rebinding time changes with the conditions varied in the experiment, we make physical interpretations of the rates between states. The rates *k*_near_ and *k*_far_ depend on how the cargo moves rotationally. We model this as *k*_near_ = *aD* and *k*_far_ = *bD*, i.e., the timescale of rotational movement is ∝ 1*/D*, where *D* depends on both size and viscosity. For a free sphere rotationally diffusing in a simple fluid, the coefficients *a* and *b* are known. However, the current setup could be complicated by the presence of the surface [19] and the microtubule. To let our model act as a simple description of this system and test whether it can capture the data, we do not a priori specify *a* and *b*, and instead let the model identify them as described below.

The rate *k*_on_ represents the binding rate of the motor to the microtubule, assuming it is close enough to reach. On rates can be thought of as having two different regimes in opposite extremes: a diffusion limited regime, where the proteins spend most of their time far from each other and bind quickly once they come near, and a reaction limited regime, where the proteins come near each other many times before they successfully bind [20]. In the diffusion limited case, changing the viscosity of the solution in which the proteins are diffusing will affect the average binding rate. In the reaction limited regime, changing the viscosity of the solution is predicted to have no effect on average binding time. Since little is known about how kinesin motors explore space and bind to a microtubule, we do not know if this binding rate should be closer to diffusion limited or reaction limited a priori. Therefore we construct two different models. The first, assuming motor binding is in the reaction limited regime, yields *k*_on_ independent of viscosity of the fluid. The second, assuming motor binding is diffusion limited, yields *k*_on_ ∝ 1/*η*. Either way, we assume *k*_on_ does not depend on the size of the cargo. We perform the subsequent analysis under both models. In the main text, we consider the model where *k*_on_ does not depend on *D*. An analogous investigation for the diffusion limited model is shown in S4, where we surprisingly find that our main conclusion and quantitative estimates are the same.

We solve the model for the distribution of times to get to the engaged state. The model has an additional unknown, *f*, describing the probabilistic mixture of cargos starting in the far state versus in the near state. From this solution, we derive the mean time to the engaged state, which has a compact form:

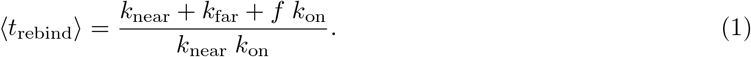

We next consider the limit as *D* becomes large. This puts binding in a regime where the motor visits the microtubule many times before binding occurs. In this regime, the binding time should be inversely proportional to the ratio of the time the cargo spends in the near state and the far state, which is itself the same as the ratio between the areas on the cargo in which a motor can be located where it is able to bind vs. unable to bind. We term this ratio *r*, and are able to find it as a function of the cargo size using simulation. Further description is given in supplemental figure S3. Enforcing this on the mean rebinding time eliminates one of the unknown coefficients, leaving only *a*, which does not depend on cargo or environment properties. Since *r* is determined by bead and motor geometry and *D* is set by cargo size and environment viscosity, we are left with unknown parameters *f*, *a*, and *k*_on_. Substituting our physical assumptions into 1 yields

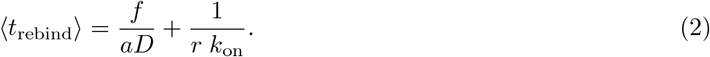

In figure 3b, we show that this model is capable of producing rebinding time distributions with a variety of qualitative behaviors. It can produce exponential-like distributions for some parameter values, and bi-exponential like distributions for others. Importantly, it is able to closely match the bi-exponential distribution fit to the experiment.

We then wish to examine if the model behaves in a similar way to the data as a function of the diffusion coefficient. In figure 3c, we show that the model can exhibit different behaviors depending on *f*, the proportion of cargos that start in the far state. When *f* = 1, the model gives average behavior reminiscent of the data, with the time decreasing as diffusion coefficient increases. When *f* = 0, however, the model gives an increasing rebinding time as diffusion coefficient increases. This behavior matches expectations: if the motor starts far away from the microtubule, a lower diffusion coefficient would slow the motor’s approach to it, resulting in increasing rebinding time. When the motor starts near the microtubule, decreasing the diffusion coefficient makes it less likely to leave the the near state before it rebinds, with leaving the state becoming impossible when *D* = 0. The median rebinding time matches that of an exponential distribution with mean *k*_on_, as expected. In figure 3C and 3D we show how the each of these parameters affect the mean rebinding time as a function of rotational diffusion coefficient when other parameters are held constant.

### 2.4 Data constrains the model to yield an estimate for motor on rate

The equation has two parameters which can determined from experimental rebinding times: *f*/*a* and *k*_on_. The degrees of freedom provided by these two parameters suggest that the data will constrain them strongly. To estimate parameters of the model from the data, we use two approaches (see methods section 4.8): First, a maximum likelihood parameters for the model, and then a Bayesian approach that allows us to incorporate previous estimates of motor on rate for kinesin into our analysis in the form of a prior.

We find a single peak for the likelihood near *f*/*a* = 1, *k*_on_ =100 s^−1^. We construct a marginal likelihood for *k*_on_ to find a distribution with mean 98 s^−1^ and standard deviation 8 s^−1^. These distributions are shown in figure 4A. A plot of the model evaluated at the maximum likelihood parameters is shown in figure 4B.

**Figure 4:**
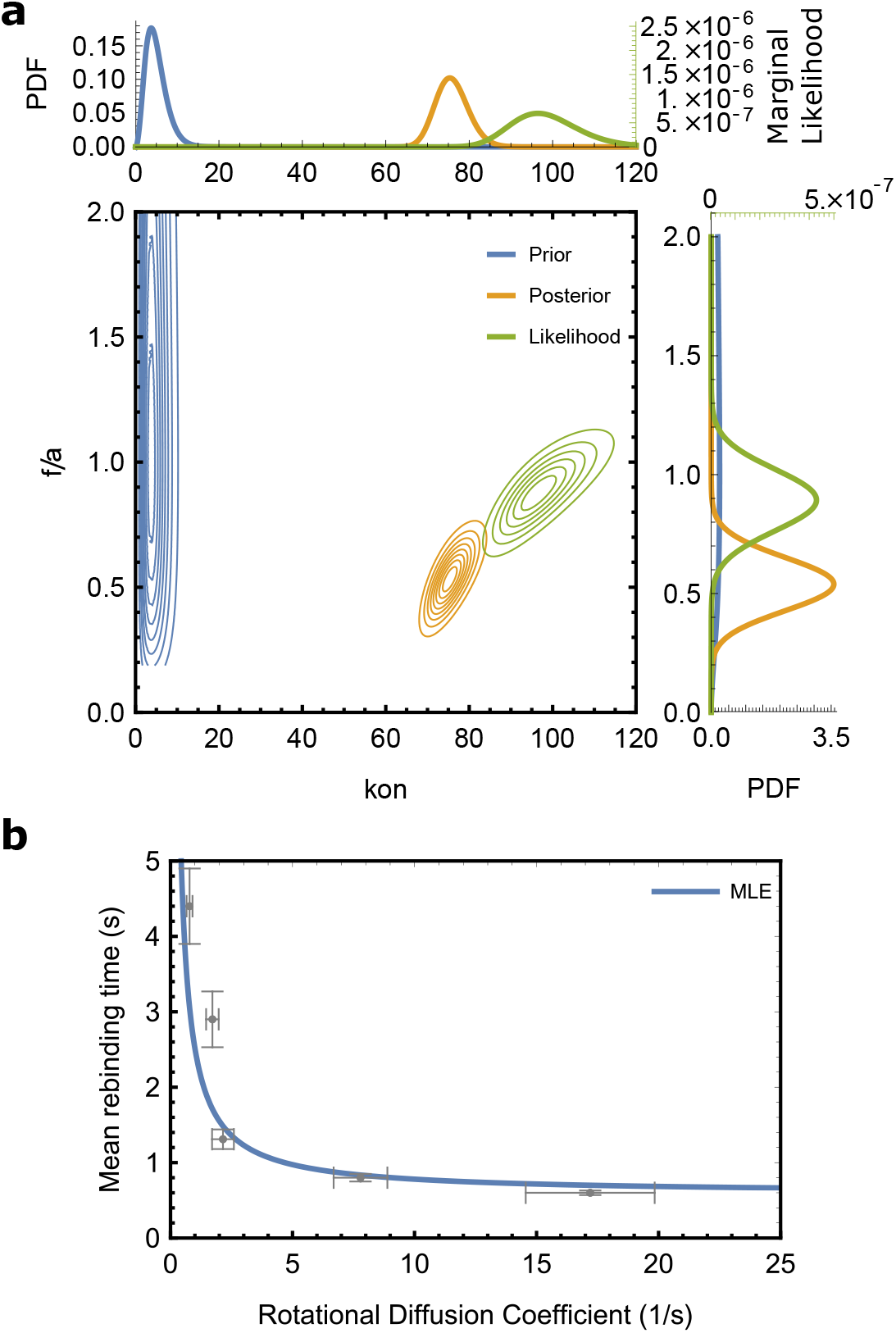
Parameter estimation for model parameters *k*_on_ and *f*/*a*. We compute the likelihood of the model parameters given the data, shown in green in (A). Contours of the joint likelihood are show in the central panel, while marginal likelihoods for each parameter are shown on above and below. Values are given on the green axis. We construct a joint prior distribution from kon values estimated in the literature and an uninformative lognormal distribution in *f*/*a*. We then compute the posterior distribution from the prior and the likelihood, yielding the joint distribution shown with contours shown in orange. The marginal distributions corresponding to these joint PDFs are shown in the top and right panels. In (B) we show the data with the model at the maximum likelihood parameters.

The parameters we find from analyzing the data for several viscosities and bead diameters in this section suggests that cargo attachment is a result of slow cargo motion combined with fast local motor binding. Further supporting this conclusion is the biexponential distribution of rebinding times found for experimental cargos (figure 1). The model parameters (*k*_on_ = 100, *f* = .32, *a* = .32) shown in figure 3b.

To incorporate previous estimates for motor on rate into our analysis, we construct a prior. As discussed in the Introduction, previous estimates for kinesin on-rate were in the range of 0.7 s^−1^ [9] to 10 s^−1^ [16], with typical values around 5 s^−1^. To summarize these findings, we construct a prior with most of the weight near 5 s^−1^, and some weight near 1 s^−1^ and 10 s^−1^. Because motor on rate must be greater than 0, we choose a Gamma distribution with mean 5 s^−1^ and standard deviation 2.5 s^−1^. We choose a LogNormal distribution for *f*/*a* with a peak at 1 and a wide spread. This prior distribution is shown in figure 4A. From the likelihood and the prior we derive the posterior probability for motor on rate. We find a distribution with mean 76 s^−1^ and standard deviation 4 s^−1^. This result indicates that the data strongly determines motor on rate in these rebinding experiments to be much higher than previous estimates.

As mentioned above, we also analyze a model in which motor binding is assumed to be diffusion limited. Surprisingly, this yields a similar *k*_on_ at water viscosity, 86 s^−1^ ± 6 s^−1^ (mean ± standard deviation) (see figure S4). To determine if the diffusion limited model explains the data better than the reaction limited model, we perform Bayesian model comparison. We find that a Bayes factor of 3.8, which we interpret as weak evidence in favor of the reaction limited model (as opposed to positive or strong evidence, indicated by Bayes factors 10-100 times higher [21]).

## 3 Discussion

While there are many established experimental and theoretical approaches to understand molecular detachment and mobility, the attachment process is less studied, and poorly understood. In this work, we provide a quantitative study of cargo-bound kinesin-1 attaching to a nearby microtubule. We find that attachment is determined by a combination of cargo rotation, which in turn depends on cargo properties and environment properties, together with the kinesin-1 attachment rate, which we estimate as *k*_on_ = 100 s^−1^.

A first guess at the motor on rate could be made by simply inverting the raw rebinding time, which yields 1.25 s^−1^ for 560 nm diameter cargos in water. Why do we infer *k*_on_ to be so much higher? Heuristically, this conclusion follows from the inevitability that if the motor is away from the microtubule, it takes for the cargo to rotate so that the motor comes near it. Even for an infinitely high motor on rate, the mean time for the cargo to bind is 0.1 s to 1 s if the motor starts far away [22]. This effect is exacerbated the fact that the motor is small in comparison to the cargo. Because of this, there are relatively few rotational orientations of the cargo which allow the motor to reach the microtubule (2.5 % to 5.3 % for our cargo sizes, see supplemental figure S3). This slows down binding in two ways: first by making visiting the region rare as the cargo rotates, and second by making the time which the cargo spends in orientations where the motor can bind short (about 0.005 s for a cargo in water, see 4.9). The few acceptable orientations and short timescale for leaving those orientations give context to why the *k*_on_ we infer is high.

Our model is general and makes few assumptions, allowing the parameter inference process to learn several features of the model from the data. We make no assumption about where on the cargo the motor begins the rebinding process, and allow the model to learn the parameter *f* from the data. We make no assumption about how much slower cargo rotational movement may be compared to free movement in water, but learn the parameter *a* from the data. We enforce on the model that rotation becomes slower as *D* decreases. The functional form of that decrease with cargo diameter and fluid viscosity hold when fluid interactions with the nearby coverslip are considered [23]. We consider both extremes of on rate dependance on viscosity and come to the similar inferences about *k*_on_. The breadth of these possibilities gives us confidence that our parameter estimates are robust, given the current understanding of the bead-motor-trap system.

Nonetheless, there could be yet-to-be-discovered confounding effects. For instance, there could be a weak association between the motor and the microtubule during the time interval when we consider the motor to be detached, with the motor heads sliding back along the microtubule and keeping the motor close. There is no evidence for this (see, e.g., [14]). In that scenario, a much lower transition rate between the sliding state and the actively walking state could then account for the short rebinding time.

The on rate we find is much higher than previous estimates, introducing a potential puzzle. If naïvely plugged into a model for the run length of a two motor construct, an on rate of 100 s^−1^ predicts extremely long run length for two motors, which is not what is seen in experiments done with these constructs [4, 8–16]. In those constructs, the rate was measured for motor attachment in a cargo carrying multiple motors, once the first motor was already attached. This is *a priori* different from the attachment rate reported here, which is the first motor to bind upon cargo encounter with the microtubule. There are several possible reasons that the on-rate for the first motor might be different from the on-rate for subsequent motors. There could be direct interaction between motors, or crowding and competition for sites on the microtubule lattice. The way that the motor heads explore space when they are not bound to the microtubule, which is different between motor constructs and how they are attached into multi-motor constructs, could also drive drastic differences between effective on rates for a given scenario. One could imagine that microtubule post-translational state could also affect this exploration.

In vivo, several factors could modify the attachment process. Viscosity is perhaps 200-fold higher than water [24]. Our results suggest that a severely limiting timescale for intracellular transport is the timescale for reorganization of the cargo necessary to allow the motor access to the microtubule. Even with an infinitely high motor on rate, the time for the cargo to bind is long, especially at high viscosities [22]. To overcome this, we hypothesize that many biological cargos have surfaces with liquid properties, like lipid droplets or vesicles. This would allow motors freedom to diffuse on the two-dimensional cargo surface, unlike in the bead experiments here, and more like [25]. In [22], we use computational simulation to demonstrate that this motor freedom allows motors to find the microtubule quickly, overcoming the slow cargo rotation times observed here. Alternatively or in coordination, MAPs may tether the cargo in place or otherwise spatially constrain the motors.

## 4 Methods

### 4.1 Bead Assay

Optical trap was assembled on an inverted Nikon microscope using a 980 nm, single mode, fiber coupled diode laser (EM4 Inc). Laser power was set to achieve the trap stiffness, ktrap, of ~0.045 pN nm^−1^ for all the polystyrene beads utilized (0.43 μm, 0.56 μm and 0.89 μm).

Motor dilutions and experiments were carried out in the motility buffer (80 mm Pipes pH 6.9, 50 mm CH3CO2K, 4mm MgSO4, 1 mm DTT, 1 mm EGTA, 10 μm taxol, 1 mg mL^−1^ casein). Viscosity of the motility buffer was altered (2x, 5x and 10x water) by adding appropriate amounts of sucrose. In all the experiments, single motor kinesin coated polystyrene beads were prepared just before the measurements. K-560-His (Kinesin-1, aa 1-560; His tag at c-term) diluted to ~20 nm was mixed with ~1 pm of penta-His-biotin antibody conjugated streptavidin polystyrene beads (0.43 μm or 0.56 μm or 0.89 μm diameter, Spherotech). This ratio produced the bead binding fraction of 10-15% and was maintained to maximize the probability of finding single motor beads in the solution. The bead-motor incubation (~30 μL volume) was carried out at room temperature for 10 min. At the end of incubation, sample chamber with preassembled microtubules was washed with ~30 μL of warm filtered buffer just before injecting the incubated mixture. Experiments were carried out at RT in motility buffer supplemented with 2 mm ATP and oxygen-scavenging system (0.25 mg mL^−1^ glucose oxidase, 30 μg mL^−1^ catalase, 4.5 mg mL^−1^ glucose).

In general, small dust or debris in the solution gets pulled into the trap along with the bead. Trapped dust interferes with motor rebinding to MTs. In our assays, dust particles and aggregates of casein in the buffer were removed using a 100 kD centrifugal filter (Millipore). Another potential issue is the stage drift during measurement and it was minimized with an automated drift correction system using a xyz piezo stage (PI) and custom software.

### 4.2 Data acquisition and analysis to extract rebinding times

Activity of the kinesin motors was observed as bead displacements from the trap center in DIC video microscopy (30 fps). The binding events were also registered in photosensitive detector (PSD) at the back focal plane of the condenser. Both these data were digitized and stored into computer hard disk using data acquisition boards and custom LabVIEW software.

Matlab code was developed to extract the duration of binding events and rebinding time between the events from the high resolution PSD signal (3 kHz). Raw data was smoothed using an FFT filter (20 point) before the analysis. The kinesin binding events appear as peaks in the PSD data. The criteria used for scoring a binding event is a displacement >=20 nm from the baseline for at least 0.005 s.

### 4.3 Description of Data and Uncertainty Estimation

Our data consists of many individual rebinding times. An individual bead gives about 10 rebinding times, and about 10 different beads are measured for a given condition of bead diameter and sucrose concentration. Multiple beads come from a single prepared slide.

We group all rebinding times from beads in a given condition to derive the distribution of rebinding times for that condition. We calculate the mean and standard error of the mean for these data.

Beads are variable in size, and experimental preparations are subject to variation in sucrose concentration and temperature. Because we do not directly measure these quantities, we estimate their value and uncertainty in the following ways.

Mean bead size is provided by the manufacturer. The manufacturer also provides a distribution of sizes for 560nm beads, from which we learn the distribution of bead sizes has a standard deviation of 95 nm. Standard error of the mean is then calculated using this standard deviation and the number of beads measured.

Viscosity of sucrose solutions have been measured [18] as functions of both concentration and temperature. We use 35 % and 45 % solutions to create 5x water and 10x water viscosity solutions respectively. We set our mean viscosity values by interpolating linearly within the tabulated data of [18] for 35 % and 45 % sucrose at 22.5 °C. We estimate an uncertainty of 5 percentage points of sucrose concentration. Since the trapping laser can create local heating near the bead [26], we estimate the temperature of the system to be between the 20 °C room temperature and 25 °C. We estimate 95 % of values fall within a range set by the mean of the four extreme values (±5 percentage points of concentration at 20 °C and 25 °C).

Viscosity of water without sucrose is also a function of temperature [27]. We linearly interpolate with the tabulated values of [27] at 22.5 °C to find our mean value, and estimate 95 % of values fall within the range set by the mean of the values at 20 °C and 25 °C.

To convert our uncertainty estimates into standard errors of the mean, we divide them by 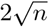, where *n* is the number of beads.

We use the the above estimates of mean and standard error for bead size (radius *R*) and viscosity (*η*) to calculate the mean and standard error for rotational diffusion coefficient, *D* = *k*_*B*_*T*/(8*πηR*^3^), with *k*_*B*_*T* =4.1 × 10^−3^ pN μm. Error was propagated using Mathematica’s ability to do automatic error propagation on values in Around[] objects.

### 4.4 Curve Fitting

Biexponential fit in figure 1 was accomplished using the fit function in Matlab 2019b. Rebinding time emperical cumulative distribution was fit to

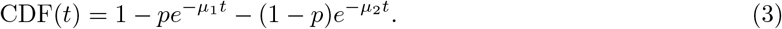

### 4.5 Cargo simulation

Simulations were done using models and code constructed previously [16, 22]. We note that unlike in [16], we include only one microtubule and one motor on the bead. Briefly, we model the bead as a sphere which moves translationally and rotationally in 3D space under forces from an optical trap, motor stepping, and diffusion. Equations of motion are given in [16]. We implement an optical trap force on the bead with the form

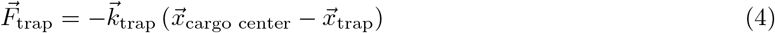

which is included in the external force term of [16]. Equations of motion take the form of stochastic ODEs, which are solved forward in time with an Eular-Maruyama scheme.

On each bead we place a single motor, which is anchored rigidly. Motors have rates of binding, unbinding and stepping which depend on the relative location of the motor anchor and microtubule.

The binding rate of the motor is

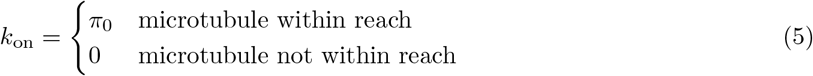

The microtubule is within reach of the motor if the distance between the location of the anchor and any point on the surface of the microtubule is less than the motor length away and the motor would not have to go through the bead to reach it. Motors are allowed to bind if the motor could reach around the surface of the bead to the surface of the microtubule. We estimate 40 nm for the length of k560 with the antibody linker.

The motor steps along the microtubule with a rate that depends on the force *F* experienced by the motor,

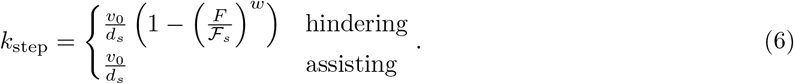

The model is based on [16], with updates to our model for motor unbinding to reflect recent findings about kinesin unbinding rate under assisting load [1]. We state the unbinding rate as

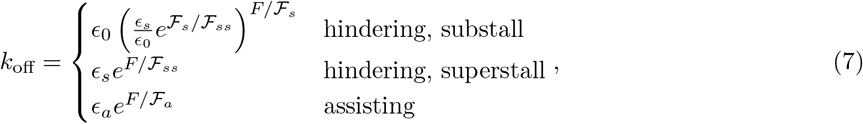

We compare with other forms that have been used in previous models in supplemental figure S1 and find that the differences are negligible.

Parameters are listed in Table 2. More detailed descriptions of rates are given in [22]. Rates are implemented through a Gillespie/Next Event method, which operates at the same time as the Euler-Maruyama implementation of the stochastic ODE solving for the equations of motion. See [16] for more details.

**Table 2:** Parameters used for detailed computational model.

| Parameter | Description | Value |
| --- | --- | --- |
| $\vec{k}_{\text{trap}}$ | Optical trap stiffness (x,y,z directions) | (.04,.04,.013)pN nm<sup>-1</sup> |
| $\pi_0$ | motor on rate | varied in simulations |
| $v_0$ | 0 force velocity | 0.75 $\mu\text{m s}^{-1}$ |
| $d_s$ | step distance | 8.2 nm |
| $\mathcal{F}_s$ | stall force | 6.1 pN |
| $w$ | force-velocity curvature index | 1.8 |
| $\epsilon_0$ | 0 force unbinding rate | 0.8 s<sup>-1</sup> |
| $\epsilon_s$ | superstall base unbinding rate | 1.6 s<sup>-1</sup> |
| $\mathcal{F}_{ss}$ | characteristic superstall detachment force | 7.6 pN |
| $\epsilon_a$ | assisting base unbinding rate | 7.4 s<sup>-1</sup> |
| $\mathcal{F}_a$ | characteristic assisting detachment force | 12.5 pN |

### 4.6 Derivation of rebinding time distribution and mean

Differential equations

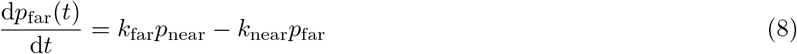

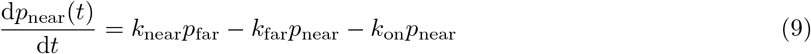

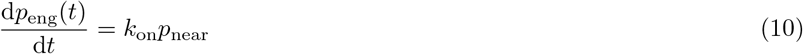

were entered into Wolfram Mathematica 12.0.0.0 and solved analytically using DSolve with initial conditions *p*_far_(0) = *f*, *p*_near_(0) = 1 − *f*, *p*_eng_(0) = 0. Binding time cumulative distribution is the same as *p*_eng_(*t*). Mean time to bind,

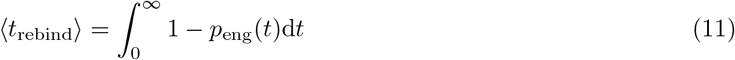

was found using Integrate. Medians were found by solving *p*_eng_ = .5 using FindRoot.

### 4.7 Determination of the area ratio parameter *r*

The area ratio is introduced in figure S3. To find the max reach locations, we began 500 simulations with a single motor located at the north pole of the cargo, positioned directly above a microtubule. We allowed the cargo to diffuse rotationally (but not translationally) until the first time at which the motor was able to bind to the microtubule. Motors were determined to be able to bind if the shortest distance around the surface of the cargo to the microtubule is less than the motor length (for more, see detailed description of simulation in [22]. We recorded this location for each simulation and took the mean to find the mean reach elevation. This process was repeated for each cargo diameter. To the data on mean reach elevation as a function of cargo diameter data we fit the function 1/ log (*ad^b^* + *e*^1/*π*^), and find *a* = 28, *b* = 1.6 with the function fit in Matlab 2019b.

### 4.8 Parameter Estimation

We fit our model to data points from each of our five experimental conditions. Each condition *i* has a mean estimated rotational diffusion coefficient *μ_D,i_* and a mean measured rebinding time *μ_t,i_*. We also have the standard errors of the mean associated with each, *σ_D,i_* and *σ_t,i_*.

The mean time to bind derived from the model is stated in equation 2. Define *t*_model_(*D*, Θ) for the mean time to bind predicted by the model. This is a function of the diffusion coefficient *D* and other parameters Θ, representing *f*/*a*, and *k*_on_ for the binding rate limited model and 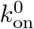 for the diffusion-limited model.

The likelihood of observing a data point (*μ_D,i_, μ_t,i_*) given our model *t*_model_(*D,* Θ) as

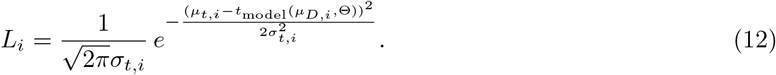

Then the likelihood of observing all the data is 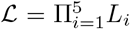. We use this likelihood 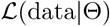 along with our prior on the parameter values *P* (Θ) to find the posterior distribution

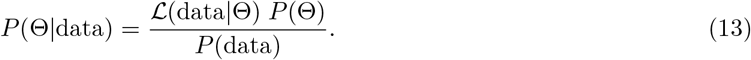

The denominator *P* (data) is determined by the condition that ∫_*S*_ *P* (Θ|data) = 1, where *S* is the domain of all possible values of parameters Θ. Numerically, we complete this integral by examining the likelihood, prior and posterior and finding limits outside of which density is negligible, then integrating over the remaining space. We find ranges of (0,60) for *k*_on_ and (0,25) for *f*/*a* contain nearly all density.

We also show marginal distributions and likelihoods, where the marginal density of a parameter is the integral of the density over all other parameters. Numerically, we calculate the marginal density by separating the space of the parameter into bins and integrating over strips. We determine *δ* by taking the range from the previous paragraph and separating it into 200 bins, which we find to be sufficient for the marginal densities to be well resolved.

### 4.9 Mean first passage time out of a spherical cap by cargo rotation

We estimate the time the cargo would spend in the near state if it were freely rotationally diffusing in water. The mean first passage time for a point on a sphere to leave a spherical cap of extent *θ* by rotation is

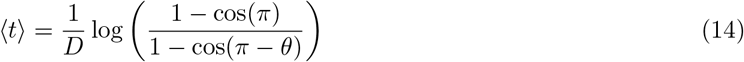

as given in [28]. By plugging in the *θ* value taken from the analysis in figure S3, the mean first passage time for a 45 nm motor starting at the south pole of a 560 nm diameter cargo in water is 0.005 s.

## 5 Acknowledgments

This work was supported by NIH R01 GM123068 to JA and SG, NIH T32 Training Grant EB009418-07 to Arthur Lander and Qing Nie, the UCI Center for Complex Biological Systems, the BEST IGERT program funded by the National Science Foundation DGE-1144901, NSF grant DMS 1763272 and a grant from the Simons Foundation (594598, QN).

## 6 Author contributions

MB, BR, SG and JA designed the study. BR performed experiments and raw data analysis. MB developed the model and simulation, and performed computational data analysis. SG and JA supervised the research.

## 7 Additional Information

The authors declare no competing interests.

---

**Figure S1:**
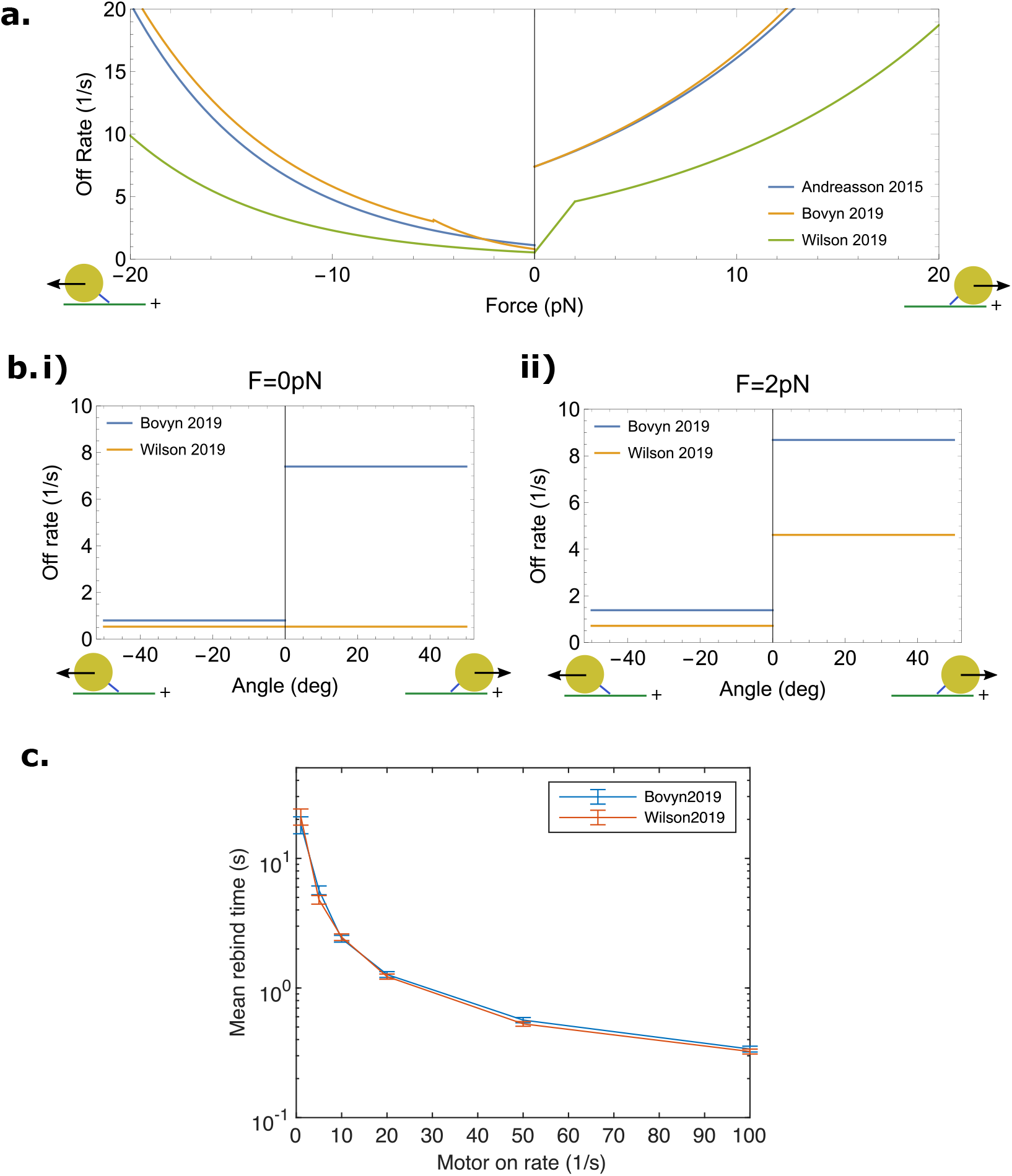
Dependance of rebinding time on assisting load off rate model In [1], off rate of kinesin is measured under hindering and assisting loads. A model with a discontinuity at 0 pN is fit to the data, as shown in a. This discontinuity is tricky, because it represents a change in loading condition from loaded with kinesin at an angle of −60° to 60°. What happens at intermediate load angles is unclear. Several models for unbinding near this continuity have been made [3, 22, 29]. In [22], the discontinuity is kept explicitly in the loading model. It is implemented as a discontinuity in angle at 0°, as shown in bi. In [29], a linear approximation is made between 0 pN load and 2 pN load (a), which can also be thought of as a off rate at 0 pN which does not change with angle (bi), but has a discontinuity at non-zero loads (bii). Because many of the binding events which are seen in the simulation but missed in the detection algorithm involve loads in this intermediate zone, the unbinding model could potentially impact rebinding times. To test this, we implemented the unbinding model from [29] into our simulation and compared mean rebinding times. As shown in c, we find no difference in rebinding times between the two models.

**Figure S2:**
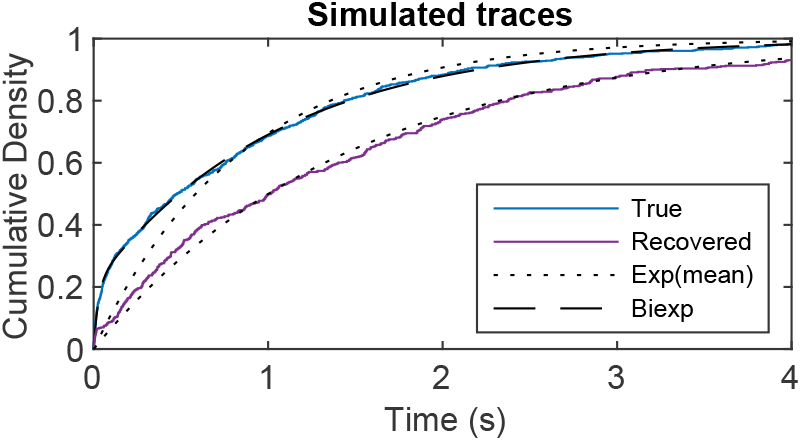
Rebinding distributions from the simulation. Empirical cumulative distributions of rebinding times, taken from the true known internal state of the simulation and recovered by the kick detection algorithm. Shown overtop are exponential distributions with the same mean as each.

**Figure S3:**
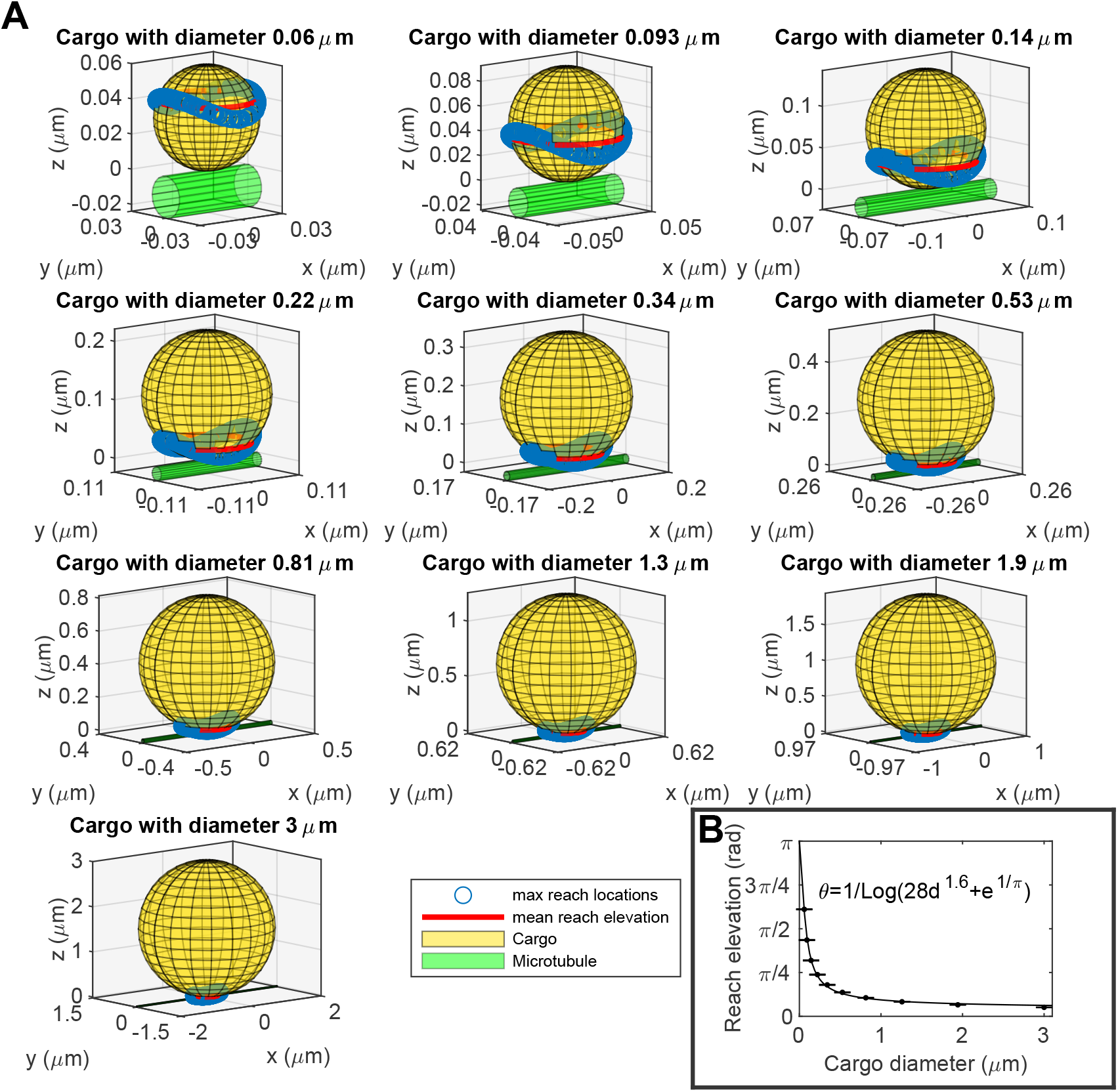
Bead and motor geometry determine area ratio In our compartmental model we introduce the parameter *r*, a ratio of areas on the cargo surface. The numerator of this ratio is the area on the cargo surface to which a motor can be bound which allows the motor to reach the microtubule. The denominator is the full surface area of the cargo. We use our simulation to determine this ratio as a function of the cargo diameter. In A, we show cargos of various sizes next to a microtubule, along with locations where motors were just barely able to reach the microtubule (blue circles). We take the mean elevation of these locations to find the mean reach elevation, shown in red. Mean reach elevation is shown as a function of cargo diameter in B. Error bars are standard error of the mean. The function *θ* = 1/ log 28*d*^1.6^ + *e*^1/*π*^ is a good fit to this data. By plugging this in to the ratio of the area of a spherical cap to the total area of a sphere, (1/2)(1 − cos(*θ*)), we arrive at the area ratio as a function of cargo diameter.

**Figure S4:**
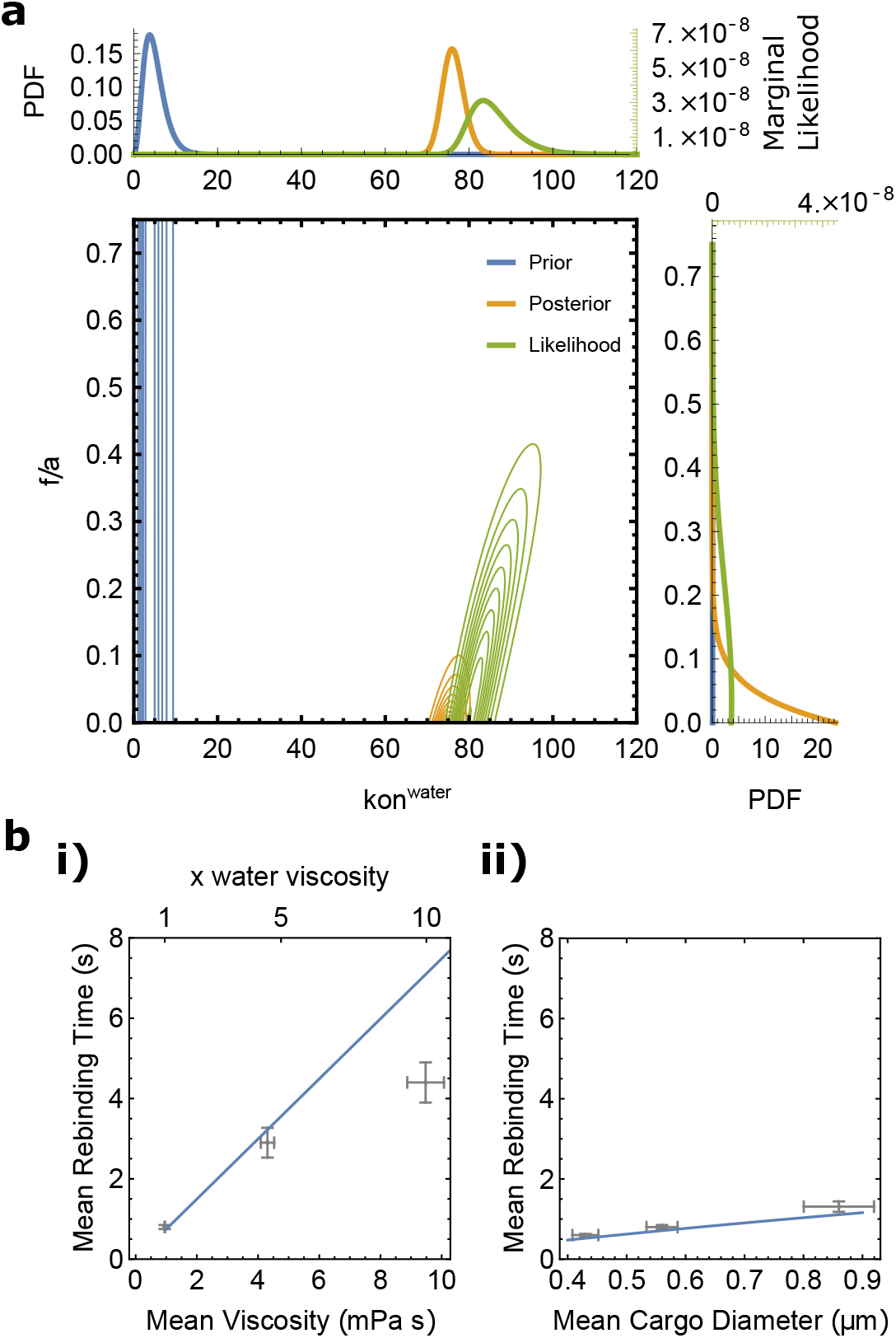
Parameter estimation for diffusion limited *k*_on_ model Analogous to equation 2, we derive an equation for the mean time to bind with motor on rate assumed to be in the diffusion limited regime, as opposed to the assumption of a reaction limited on rate. Unlike the reaction limited case, we must state the model as a function of diameter and viscosity explicitly. We find 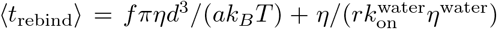. We perform the same parameter estimation on this model that was performed for the reaction limited model, and find the likelihood and posterior distributions shown in **a**. We show the data and the model with the maximum likelihood estimate parameters in **b**, as a function of viscosity in **i** and as a function of diameter in **ii**.

